# Deficiency in MICOS component *Chchd3* Compromises *Drosophila* Heart Function via mitophagy, ROS and ER Stress

**DOI:** 10.64898/2026.08.14.744045

**Authors:** Cristiana Dondi, Shuchao Ge, James Marchant, K’Leigh Guillotte, Karen Ocorr, Georg Vogler, Rolf Bodmer

## Abstract

A pair of paralogs, Chchd3 and Chchd6, two components of mitochondrial contact site and cristae organizing system (MICOS), have been identified to be candidate pathogenetic genes in congenital heart disease (CHD). Previous research found that knockdown (KD) of the single *Chchd3/6* (*Chchd3)* gene and other MICOS components in *Drosophila* impaired heart function, likely due to a deficit in mitochondrial organization, ATP production, actomyosin levels, and thus severely diminished contractility. However, the underlying mechanisms of how MICOS deficiency leads to these defects are not clear. Here, we performed genetic manipulations in the *Drosophila* heart to probe for possible interactions between MICOS-compromised mitochondria and other organelles and processes. We found that moderate reduction in *Pink1/parkin*-mediated mitophagy synergistically aggravated cardiac *Chchd3* KD phenotypes, indicating a major interaction. Further, *Chchd3* KD increased the level of reactive oxygen species (ROS) and endoplasmic reticulum (ER) stress. Interestingly, KD of *catalase* (*CAT*) also elevated cardiac ROS levels, but surprisingly did not compromise contractility either by itself or in combination with *Chchd3* KD to aggravate the cardiac phenotype. However, *CAT* overexpression (OE) in *Chchd3* KD hearts restored contractility, but only partially, even though elevated ROS due to *Chchd3* KD was fully normalized. Similarly, counteracting ER stress by overexpressing *Xbp1* (or spliced mouse *Xbp1*) also partially rescued the heart function defects induced by *Chchd3* KD. Overall, these data indicate a critical role of mitophagy and ER/oxidative stress in cardiac homeostasis involving *Chchd3*, which suggests that deficiency of MICOS function contributes to heart dysfunction via multiple stress responsive pathways.

## Introduction

The mitochondrial contact site and cristae organizing system (MICOS) is crucial for maintaining the architectural integrity and physiological functions of mitochondria [1]. Located at the edge of the mitochondrial cristae (**Fig. 1**), they serve as a distribution center for electron transport chain components into the cristae and are involved in diverse cellular functions [2]. Rare, predicted-damaging genetic variations of Chchd3/Mic19, Chchd6/Mic25 and other MICOS components have been identified in human patients with hypoplastic left heart syndrome (HLHS), a deadly congenital heart disease (CHD), which suggests that MICOS components are candidate genes in CHD pathogenesis [3]. Furthermore, splicing variants of Chchd3 were detected in a zebrafish model of HLHS, further supporting involvement of Chchd3 in heart development and function [4]. Moreover, in aging mice, the expression of Chchd3 and Chchd6 declines in both cardiac and skeletal muscles, consistent with the dystrophy and declining performance of muscles with age [5, 6]. *Drosophila* has only one ortholog of the paralogous genes (*Chchd3*), and in loss-of-function studies *Chchd3* cardiac knockdown (KD) reduced the abundance of sarcomeric actin and myosin in cardiomyocytes and much diminished heart contractility, which is accompanied by reduced ATP production [3].

**Figure 1:**
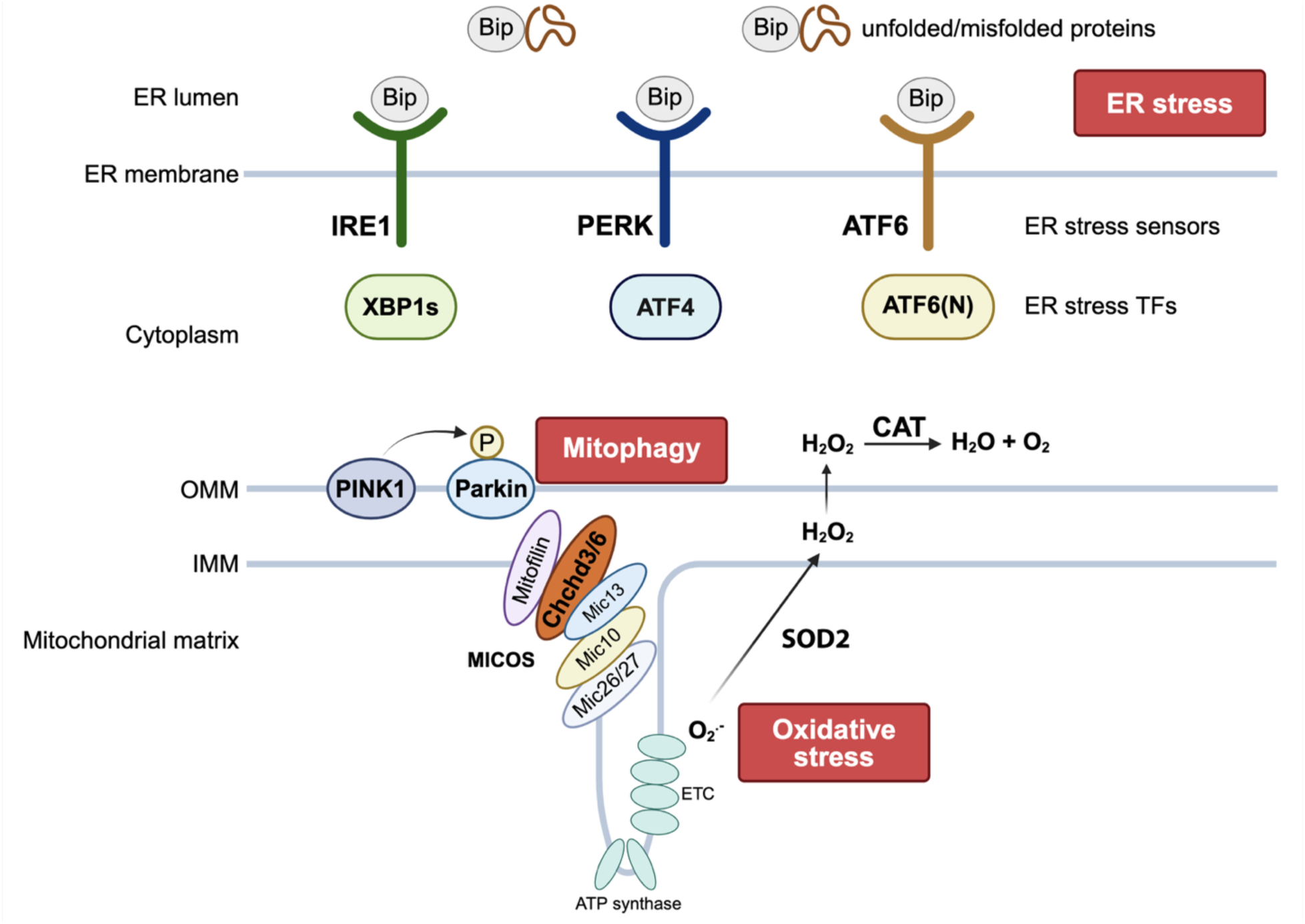
(Top) Diagram depicting mitophagy, oxidative stress and ER stress pathways. (Bottom) Diagram depicting MICOS complex genes, and the production and degradation of ROS.

In our previous study [3], we found that compromised mitochondrial fission/fusion can contribute to the observed heart damage inflicted by cardiac Chchd3 KD. Using genetic interaction studies, reducing *Drp1* function in the *Drosophila* heart synergistically aggravated the heart defects due to moderate *Chchd3* KD [3], suggesting the involvement of mitochondrial quality controls. Mitochondrial homeostasis also depends on mitophagy mechanisms regulated by the *PINK1* kinase and *parkin* E3-ligase genes, whose products accumulate on malfunctioning mitochondria to target them for degradation, including in muscles [7].

Other pathways may also contribute to the cardiac pathology due to compromised MICOS function. For example, cellular stresses often produce reactive oxygen species (ROS) as a byproduct of metabolic processes, such as OXPHOS in the electron transport chain (ETC) in mitochondria as well as by NADPH oxidases [8]. In physiological condition, ROS is converted to chemically stable molecules by superoxide dismutases (SODs) and catalase (CAT) [9] (**Fig. 1**). Although ROS are also physiological signals [10, 11], unusually high levels of ROS are generally considered harmful to the organism, and has been reported in various stressful or pathological conditions, including aging [12], cancer [8], diabetes [13] and cardiovascular disease [14]. In addition, ER stress happens when harmful cellular and environmental stimuli lead to the accumulation of misfolded or unfolded proteins in the ER lumen [15]. To restore ER to the normal physiological condition, the unfolded protein response (UPR) pathway is activated [15]. There are three main branches of ER stress-induced UPR, each with unique stress sensors and transcription factors mediating specific transcription responses. Depending on the activated downstream genes, the cell will either attempt to restore homeostasis or undergo apoptosis [16]. ER stress is involved in multiple human pathological conditions, such as in liver [17], pancreas [18] as well as the heart [19].

In this study, we used genetic interaction studies to investigate mitophagy, oxidative stress and ER stress to determine whether they may contribute to cardiac dysfunction as a consequence of loss-of-MICOS-function. Manipulation of each pathway simultaneously with moderate (“sensitized”) *Chchd3* KD was found to synergistically aggravate or ameliorate heart dysfunction, indicating significant involvement of all three processes in regulating heart function. Among the three pathways, ER stress plays a crucial role, since overexpressing the transcription factor Xbp1 significantly restored the contractility deficit in *Chchd3* KD hearts. In sum, compromised MICOS complex function causes heart dysfunction via multiple stress pathways.

## Results

### *Pink1/parkin* KD aggravates heart dysfunction induced by *Chchd3* KD

Previously, we found that *Chchd3* KD diminished heart contractility, actomyosin levels and caused mitochondrial fission/fusion defects [3]. Since mitophagy is an essential process in clearing damaged mitochondria [20], we assessed mitophagy with a mito-QC reporter, containing a mitochondria-targeted GFP-mCherry fusion protein (see methods). Once damaged mitochondria enter the lysosome, only mCherry fluorescence is observed, as GFP is sensitive to its acidic environment [21]. Using this system, we observed a significant loss of GFP fluorescence compared to that of mCherry suggesting that cardiac *Chchd3* KD results in lysosomal degradation of damaged mitochondria (**Supp Fig. 1**). As previously reported [3], we confirmed that ATP synthase staining was also reduced in *Chchd3* KD (**Supp Fig. 1**).

To assess how mitophagy is involved in cardiac defects induced by *Chchd3* KD, we combined *Chchd3* KD with KD of *Pink1* or *parkin* (*park*), two genes that control induction of mitophagy (**Fig. 2**, [7, 22]). For this purpose, we used a moderately-strong *Chchd3* KD line (C1) in combination with the heart-specific *Hand^4.2^*-Gal4 driver [23, 24] and the *in vivo* tdtK reporter [25] to monitor heart function [26]. This ‘sensitizer’ line allows to finely tune the KD, in that strong *Chchd3*-C1 KD (at 25°C) considerably reduced contractility (Fractional Shortening, FS) and caused dilation (increased systolic and diastolic diameters - ESD, EDD) relative to controls, whereas KD at 21°C KD produced only mild defects (**Fig. 2, Supp Fig. 2**)(see also [3]).

**Figure 2.**
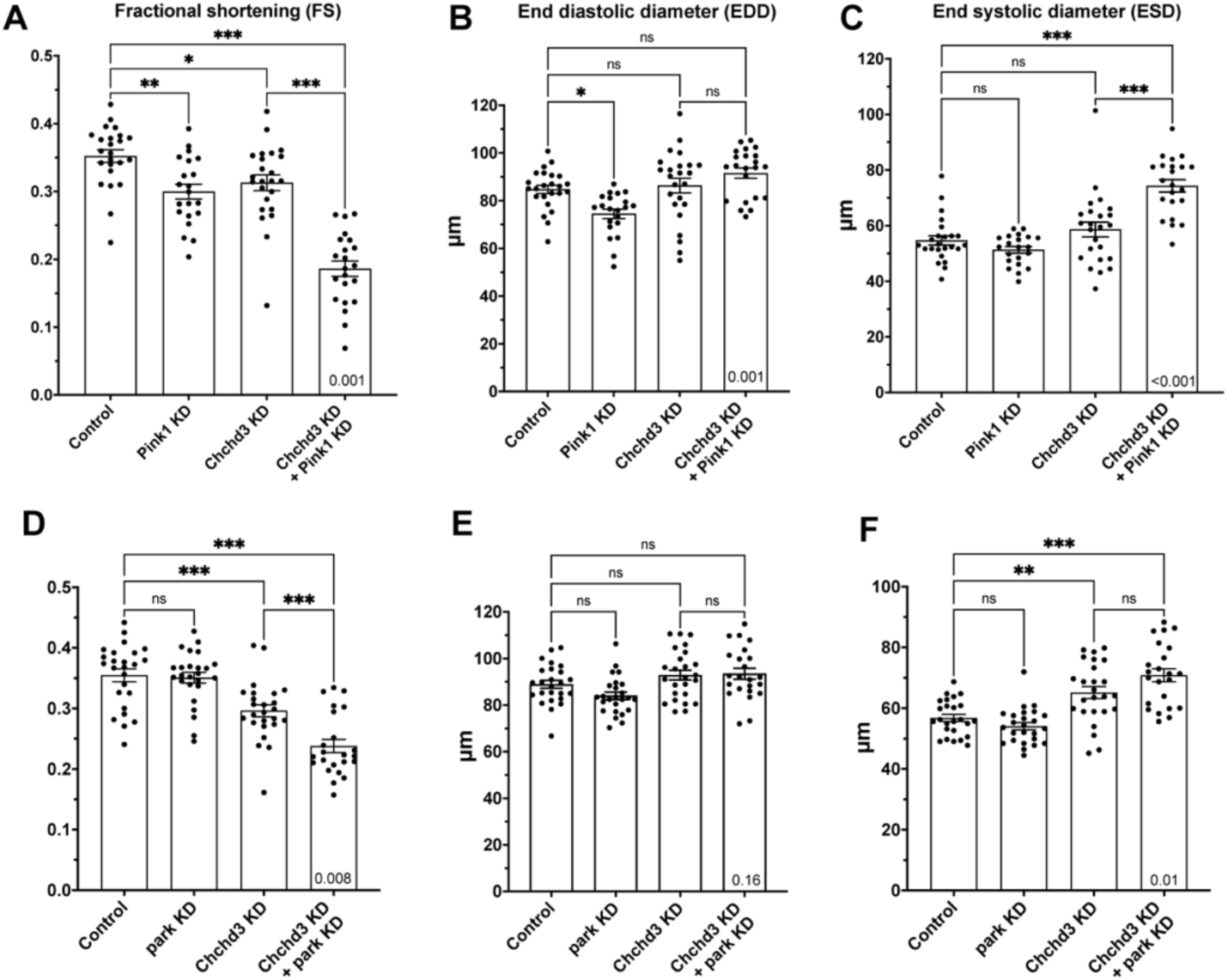
Mitophagy genes, *Pink1* and *parkin* (*park*), genetically interact with *Chchd3*. Individual KD at 21°C of *Chchd3*, *Pink1* or *park* had only mild or moderate effects on heart contractility (FS; **A,D**) or diameters (EDD, ESD; **B,C,E,F**). (**A-C**) Combined KD of *Pink1* and *Chchd3* significantly reduced FS, more than either KD alone (**A**). The double KD exhibited a trend in EDD increase (**B**), and a significant increase in ESD (**C**), thus the dramatic reduction in FS (**A**). (**D-F**) Combined KD of *park* and *Chchd3* also synergistically aggravated the reduction in FS (**D**) and increase in ESD (**F**), but EDD was less affected (**E**). FS, fractional shortening. EDD, end diastolic diameter. ESD, end systolic diameter. Bar graphs show mean with SEM. Data were analyzed by one-way ANOVA with Bonferroni multiple comparisons. *<0.05, **<0.01, ***<0.001 and ns, not significant. In each panel, inside of the last data column at the bottom the interaction p-value from two-way ANOVA analysis is indicated. Same analysis was performed in subsequent figures.

To test for synergistic genetic interactions, we crossed this C1-sensitizer line with *Pink1/park* RNAi (KD) or overexpression (OE) lines (see Methods and [3]). Individually, *Chchd3*, *Pink1* or *park* KD at 21°C caused only mildly altered FS and EDD/ESD (**Fig. 2A-F**). However, *Chchd3* KD combined with *Pink1* or *park* KD significantly increased ESD without much effect on EDD, which resulted in a dramatic decrease in FS, thus further synergistically aggravating contractile defects caused by *Chchd3*-C1 KD (**Fig. 2A-F**).

We also tested whether the strong contractility defects of *Chchd3*-C1 KD at 25°C could be rescued by elevating *Pink1/park* expression in the heart. However, cardiac-directed *Pink1* or *park* OE reduced FS, as does KD of *Chchd3*, and neither the *Pink1* nor *park* OE in combination with *Chchd3* KD resulted in any further heart defects (**Supp. Fig. 2**), although there was a significant interaction observed. This unexpected result suggests that OE of *Pink1/park* is not beneficial for mitochondrial health. Nevertheless, *Pink1/park* function does contribute to heart health, since their loss-of-function impairs contractility in combination with moderate *Chchd3*-C1 KD (at 21°C), indicating a significant genetic interaction. Overall, these findings support a genetic interaction between *Chchd3* and *Pink1/park*–mediated mitophagy but does not preclude that other stress mechanisms also contribute to *Chchd3*-dependent regulation of heart function.

### Oxidative stress participates in heart dysfunction mediated by *Chchd3* KD

While eliminating damaged mitochondria, mitophagy also removes reactive oxygen species (ROS). Given the mitochondrial defects due to *Chchd3* KD [3] and the well-established role of ROS in pathological conditions [27], we postulated that ROS level may be altered in *Chchd3* KD hearts. We examined ROS level in the fly heart with dihydroethidium (DHE), which emits red fluorescence upon oxidation by ROS [10]. Hearts with *Chchd3* KD displayed markedly increased DHE fluorescence relative to controls (**Fig. 3A, B**), indicative of substantial oxidative stress due to ROS accumulation. We wondered whether OE of the ROS-scavenging enzymes *catalase* (*CAT*) and *superoxide dismutase 2* (*SOD2*) (**Fig. 1**) can counteract ROS accumulation and *Chchd3* KD-mediated cardiac defects. Indeed, combining *Chchd3* KD with *CAT* OE fully reversed ROS accumulation (**Fig. 3C, D**), suggesting that reducing oxidative stress upon *Chchd3* KD may also lessen the heart defects. Indeed, lowering ROS levels by *CAT* or *SOD2* OE in the presence of *Chchd3* KD restored FS relative to *Chchd3* KD alone, thus resulting in a partial rescue of heart contractility (**Fig. 3E, H**). *SOD2* OE, but not *CAT* OE, also rescued *Chchd3* KD-mediated chamber dilation (EDD or ESD) (**Fig. 3F, G, I, J**).

**Figure 3.**
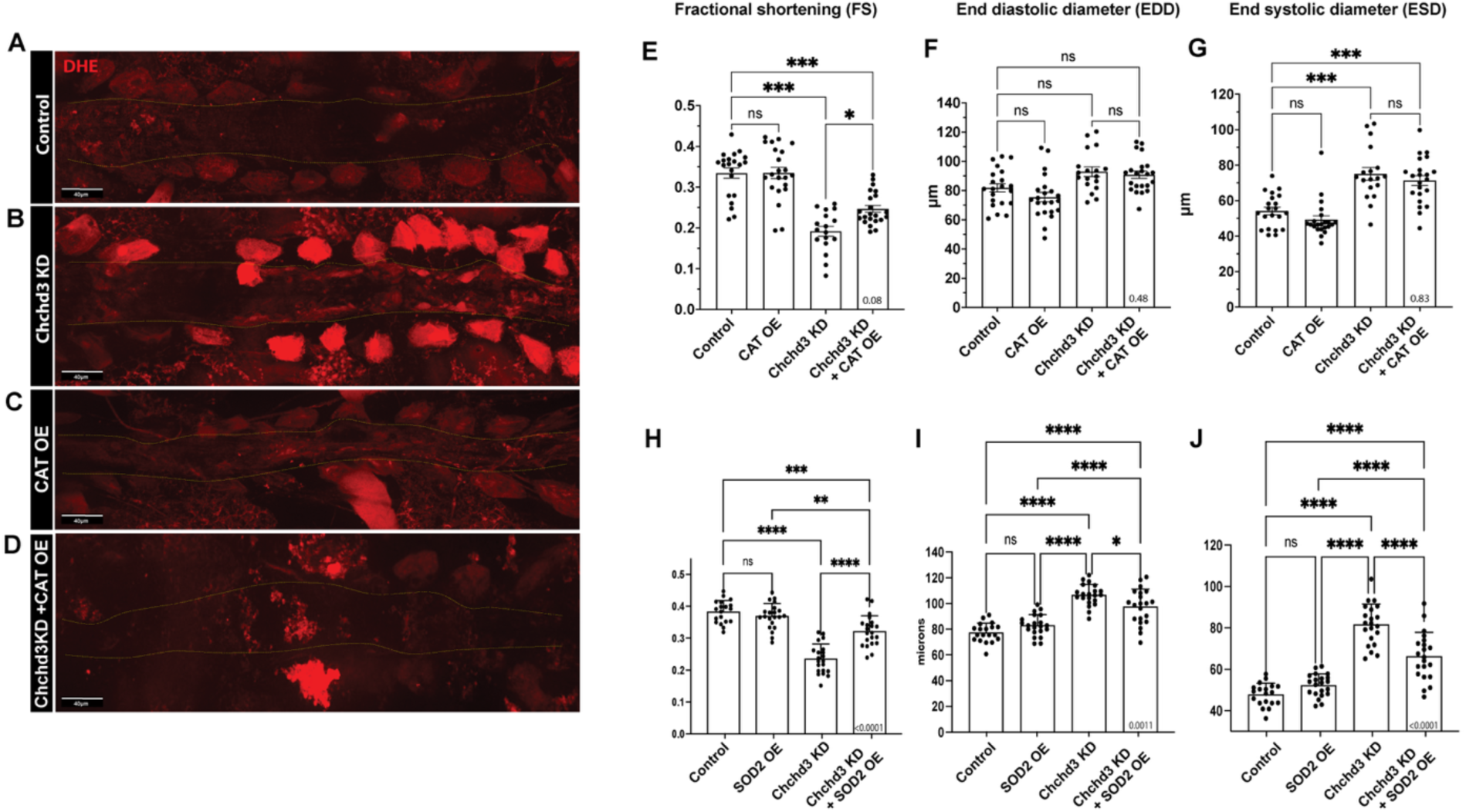
*Chchd3* KD caused oxidative stress and heart dysfunction that is partially counteracted by overexpressing ROS scavengers. (**A-D**) ROS level examined by DHE staining of flies raised at 25°C. Compared to control (**A**), *Chchd3* KD hearts (**B**), but not *CAT* OE (**C**), exhibit increased ROS levels in cardiomyocytes (inside dashed lines) and pericardial cells. However, co-expression *CAT* restored excessive ROS accumulation due to *Chchd3* KD to normal levels (**D**). (**E-J**) Reducing ROS level by *CAT* or *SOD2* OE partially rescued the functional defect inflicted by *Chchd3* KD, notably by partial restoration of contractility (**E, H**), compared to *Chchd3* KD alone. *SOD2* OE, but not *CAT* OE, also rescued chamber dilation (EDD, ESD) due to *Chchd3* KD (**I-J**). A-D, bar, 40 μm.

We also performed genetic interaction experiments similar to those conducted for mitophagy: KD at 21°C of both *CAT* and *Chchd3*-C1 sensitizer. As expected, both *Chchd3* and CAT KD moderately increased ROS levels (**Supp Fig. 3A-C**), and the combination caused a further enhancement (**Supp Fig. 3D**). In contrast, combined *Chchd3* and CAT KD did not cause a further reduction in FS relative to *Chchd3* KD alone (**Supp Fig. 3E**) but did result in additional chamber dilation (EDD/ESD), although the interaction was not synergistically significant (**Supp Fig. 3F, G**). Similarly, siRNA KD of *CHCHD3* or *CAT* in human iPSC-derived cardiomyocytes resulted in increased ROS levels **(Supp Fig. 3H, I)**.

Taken together, these results demonstrate that *Chchd3* deficiency induces oxidative stress in the heart and that enhancing ROS scavenging can partially ameliorate the associated functional defects. However, increasing ROS levels *per se*, as observed with *CAT* KD (**Supp Fig. 3C**) does not result in dramatic heart function defects, and further elevating ROS when combined with *Chchd3* KD does not enhance the defects upon *Chchd3* KD (**Supp Fig. 3E-G**). This is consistent with previous observations [10, 11] and emphasizes the conclusion that the observed functional defects, such as with *Chchd3* KD, are not exclusively due to the associated increase in ROS levels. In addition, the incomplete functional rescue by preventing ROS accumulation suggests that additional mechanisms downstream of *Chchd3* KD contribute to cardiac pathology, such as mitophagy (**Fig. 2**) and ER stress (see below).

### *Chchd3* KD induces ER stress in *Drosophila* heart

The pronounced elevation of ROS levels observed in *Chchd3* KD hearts prompted us to investigate whether ER homeostasis is also affected, as protein folding within the ER is known to be sensitive to ROS levels [28]. To determine whether ER stress was induced by *Chchd3* KD, we first measured the expression of the chaperone protein in ER stress, *BiP* (CG4147, Hsc70-3), by qRT-PCR. Chchd3 KD hearts exhibited significant upregulation of *BiP* relative to controls (**Fig. 4A**). In situ RNA detection by hybridization chain reaction (HCR) staining also showed stronger BiP expression in *Chchd3* KD heart than control (**Fig. 4B-E**). Both results indicated the induction of ER stress by *Chchd3* KD.

**Figure 4.**
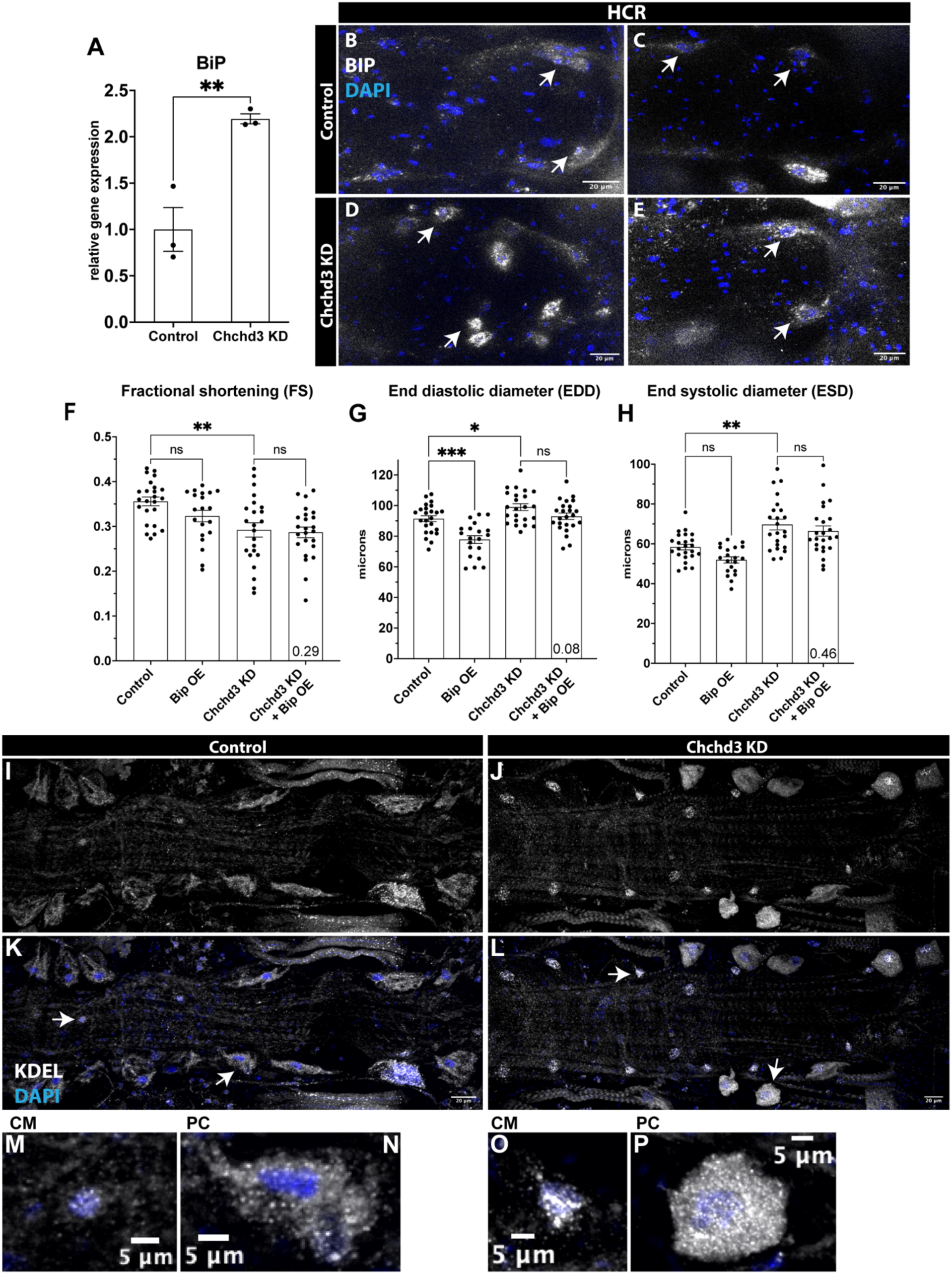
*Chchd3* KD caused ER stress. **(A)** Expression of *BiP*, an ER stress chaperon, was significantly increased in *Chchd3* KD hearts as shown in qRT-PCR and **(B-E)** *in situ* hybridization experiments (arrows show cardiomyocyte nuclei). **(F-H)** Cardiac *BiP* OE by itself caused a moderate constriction of the heart but did not rescue the reduced cardiac contractility induced by *Chchd3* KD. **(I-L)** Immunostaining against KDEL receptor demonstrated stronger fluorescence in *Chchd3* KD (**J,L**) than control (**I,K**), indicating *Chchd3* KD-induced ER stress in the heart. **(M-P)** Magnifications of cardiomyocytes and pericardial cells (arrows in **K,L**) in *Chchd3* KD (**O,P**) and control (**M,N**). **I-L**, bar, 20 μm. **M-P**, bar, 5 μm. CM, cardiomyocyte. PC, pericardial cell.

To further explore the functional relevance of ER stress, we conducted genetic interaction studies. Notably, heart-specific KD of *BiP* resulted in lethality, even at 21°C, which is likely reflecting its crucial role as a protein folding chaperone in the ER stress response. Even though *BiP* expression is increased upon *Chchd3* KD, we wondered if additional upregulation of *BiP* provided further cardiac protection and ameliorate the heart defects. However, OE of *BiP* in *Chchd3* KD hearts did not normalize the cardiac functional defects, and by itself caused moderate chamber constriction (**Fig. 4F-H**), indicating that further increasing BiP levels does not rescue the dysfunction caused by *Chchd3* KD.

We also performed immunostaining of the ER with antibodies against KDEL receptor [29], a known ER stress marker (**Fig. 4I**). Compared to the control, *Chchd3* KD hearts showed stronger KDEL receptor staining in both cardiomyocytes and pericardial cells (**Fig. 4I-P**), further supporting the presence of significant ER stress. These findings demonstrated that *Chchd3* KD induces ER stress, in both cardiomyocytes and pericardial cells of the *Drosophila* heart.

### *Xbp1* OE reduces cardiac contractility defects induced by *Chchd3* KD

We next investigated whether specific ER stress signaling pathways were involved in cardiac dysfunction caused by *Chchd3* KD, focusing on the IRE1-XBP1 branch (**Fig. 1**), the most conserved among the three UPR branches [30]. Heart-specific *Ire1* KD by itself did not alter cardiac function. However, combined KD of *Chchd3* and *Ire1* resulted in moderately reduced FS with a trend towards a significant interaction (p=0.06), as well as in chamber dilation (EDD, ESD), compared to either KD alone (**Fig. 5A-C**).

**Figure 5.**
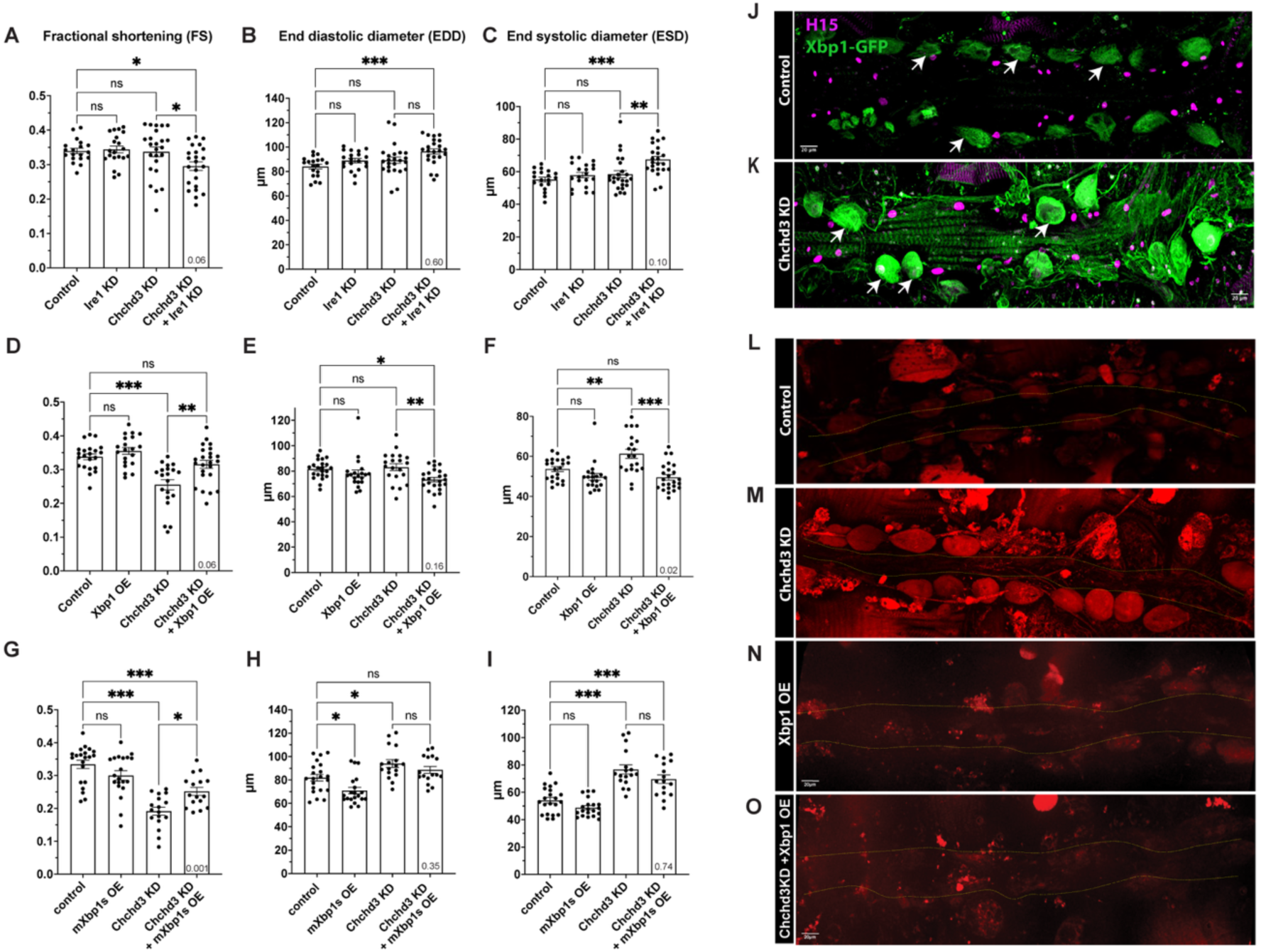
Ire1/Xbp1 branch of ER stress: *Xbp1* OE rescued contractility defects induced by *Chchd3* KD. **(A-C)** Individual KD at 21°C of *Ire1* or *Chchd3* caused little heart defects, but in combination caused a decrease in contractility (FS) and increase in diameters (especially ESD). FS exhibited a trend in genetic interaction (**A**, interaction p-value: 0.06). **(D-F)** At 25°C, OE of fly Xbp1 had no effect, and KD of *Chchd3* caused the expected dilation and decreased contractility. In combination, however, FS, EDD and ESD were restored to control levels. FS and EDD exhibited a trend in genetic interaction (**D,E**), and significant genetic interaction was observed for ESD (**F**). **(G-I)** Similarly, OE at 25°C of a spliced cDNA of mouse Xbp1 had little effect on heart function, but in combination with *Chchd3* KD significantly restored contractility (FS), with an interaction p-value of 0.001 (**G**). **(J, K)** Expression of *Xbp1*-GFP splicing sensor (see Methods) in control hearts showed GFP fluorescence weakly in cardiomyocytes (nuclei labeled with H15 antibodies) and moderately in pericardial cells (arows), indicating low levels of *Xbp1* splicing. In *Chchd3* KD hearts, however, GFP fluorescence is much increased, indicating elevated *Xbp1* splicing that suggests a robust ER stress response. **(L-O)** ROS levels were examined by DHE staining at 25°C. Compared to control and *Xbp1* OE alone (**L,N**), fluorescence was much increased upon *Chchd3* KD (**M**), indicative of the expected oxidative stress. The combination of *Chchd3* KD and *Xbp1* OE abolished DHE fluorescence, indicative of a complete protection from oxidative stress induced by *Chchd3* KD (**O**).

Upon ER stress, IRE1 branch activation causes splicing of the transcription factor XBP1 to allow its product to execute targeted downstream gene expression [31]. We first examined heart function upon *Xbp1* KD alone and in combination with *Chchd3* KD but did not observe any defects nor synergistic interaction with *Chchd3* KD (**Supp Fig. 4A-C**). Next, we asked whether *Xbp1* OE could ameliorate compromised heart function due to *Chchd3* KD. For this purpose, we overexpressed either full-length fly *Xbp1* or the spliced version of mouse *Xbp1* (*mXbp1s* [32], see methods). OE of full-length fly Xbp1 alone had little effect on cardiac parameters, but in combination with strong KD of *Chchd3* (at 25°C) caused a clear trend towards restoring contractility (FS, interaction p=0.06) and chamber dimensions (**Fig. 5D-F**). Consistently, OE of *mXbp1* robustly rescued FS (**Fig. 5G**), in this case with a highly significant interaction p-value (p=0.001). The combined effects on EDD and ESD seem to be additive rather than interactive (**Fig. 5H, I**). Together these findings indicate that *Xbp1* OE robustly rescues contractility defects of *Chchd3* KD hearts, even though *Xbp1* KD alone did not cause an appreciable synergistic aggravation (**Supp Fig. 4A-C**).

The unconventional splicing of *Xbp-1* RNA is conserved, an event for which a sensor is available in flies, *Xbp1*-EGFP [33]. Expression of this sensor in wildtype hearts exhibits weak EGFP fluorescence, indicating low levels of Xbp1 splicing. In *Chchd3* KD hearts, however, EGFP fluorescence is dramatically increased, indicative of elevated ER stress (**Fig. 5J, K**). Furthermore, given the elevated oxidative stress of *Chchd3* KD hearts (**Fig. 5L-M**), we examined whether *Xbp1* OE could also attenuate ROS levels. Indeed, DHE staining revealed that *Xbp1* OE dramatically reduced ROS to wildtype levels compared to *Chchd3* KD alone (**Fig. 5N, O**), mirroring the protective effect of *CAT* OE (**Fig. 3A-D**). The antioxidant activity of *Xbp1* OE is consistent with prior findings in mouse embryonic fibroblasts [34].

We also investigated whether the other two branches of ER stress, the UPR sensing PERK-ATF4 and ATF6 pathways (**Fig. 1**), participate in modifying *Chchd3* KD-induced cardiac dysfunction. *PERK* KD alone only mildly affected cardiac function, but combined KD of *Chchd3* and *PERK* led to somewhat worsened cardiac phenotypes that were statistically additive, not synergistic (**Supp. Fig. 5D-F**). *ATF4* or *ATF6* KD had no effect alone, and in combination did not worsen the heart defects inflicted by *Chchd3* KD (**Supp. Fig. 5D-F**).

Together, these results indicate that KD of the PERK and ATF6 ER stress components only moderately exacerbate *Chchd3* KD-induced cardiac dysfunction. In contrast, OE of *Xbp1*, the downstream effector of the IRE1 branch, was able to considerably improve *Chchd3* KD-induced cardiac contractile deficiency, highlighting the selective importance of XBP1-mediated signaling in this context.

## Discussion

In this study, we have identified mitophagy, oxidative stress, and ER stress as molecular pathways that genetically interact with *Chchd3*, a component of the MICOS complex, to regulate heart function in *Drosophila*. We have previously found that regulators of mitochondrial fission/fusion, particularly *Drp1* also interact with *Chchd3* [3]. Together, these pathways form a genetic regulatory network centered on *Chchd3* contributing crucially to the maintenance of heart function.

Using specific indicators, we observed that KD of *Chchd3* in the *Drosophila* heart resulted in increased mitochondrial degradation and elevated oxidative and ER stress. We then tested whether manipulating components of these stress-related pathways further aggravated or ameliorated the cardiac functional defects induced by *Chchd3* KD, thus probing for functional interactions of these pathways. Indeed, diminishing either of these stress pathways exacerbated the defects induced by *Chchd3* KD, while mitigating these stressors could at least partially rescue cardiac function. Our results demonstrated that deficiency in a MICOS component, *Chchd3*, leads to cardiac dysfunction through involvement of both mitophagy and ER stress mechanisms. Interestingly, mitigating oxidative stress by *CAT* OE and measured by DHE staining completely rescued ROS accumulation caused by *Chchd3* KD, but only partially rescued the heart function defects. This suggests that mechanisms other than ROS accumulation are responsible for the observed cardiac dysfunction that results from compromised MICOS function, including deficient mitophagy, ER stress and fission/fusion defects.

To investigate the mechanisms by which deficiency of the MICOS component *Chchd3* impairs heart function, we initially hypothesized that the dysfunction might originate from impaired oxidative phosphorylation (OXPHOS), given that MICOS is essential for the formation and maintenance of mitochondrial cristae [35], where OXPHOS occurs. Supporting this, our previous work demonstrated that *Chchd3* KD significantly reduced ATP production in the *Drosophila* heart [3]. Similar reduction in mitochondrial respiration was also observed in HeLa cells [36] and mouse liver [37] upon *Chchd3* KD/KO, which also resulted in decreased cristae numbers in both models [36, 37]. Consistently, an increase in ATP production was observed in mouse liver with *Chchd3* OE [38]. These studies revealed the important role of *Chchd3/MICOS* in OXPHOS.

In our current study, *Chchd3* KD also led to a marked increase in ROS, a byproduct of OXPHOS. While the precise metabolic alterations induced by *Chchd3* KD remain to be elucidated, they likely involve disruption in mitochondrial structure and electron flow. Since eliminating ROS in *Chchd3* KD hearts only partially rescued dysfunction, it is likely that other mechanisms, such as compromised mitophagy, fission/fusion and ER stress, also contribute to heart dysfunction, separate from oxidative stress. Indeed, OE of *Xbp1*, significantly ameliorated the cardiac functional defects induced by *Chchd3* KD. Again, the beneficial effect was partial. Thus, reducing ER stress parallels the benefits of ROS scavenging, suggesting that both ER and oxidative stress pathways are involved in mediating the detrimental effect of compromised MICOS function.

It is still an open question how *Chchd3* deficiency leads to impaired mitophagy and induction of ER stress. They may be consequential responses to the insufficient energy supply and accumulation of excessive ROS. Alternatively, MICOS itself may function as a structural hub facilitating interactions with molecules involved in mitophagy and ER function. Such interactions may be disrupted by compromised integrity of MICOS. In mammals, MICOS is organized into two subcomplexes: Mic60-Mic19(Chchd3)-Mic25(Chchd6) and Mic10-Mic26-Mic27, bridged by Mic13 [1, 2]. Notably, the three subunits of the Mic60 subcomplex all physically interact with SAMM50 on the outer mitochondrial membrane [36, 39]. It has been reported that SAMM50 regulates PINK1-parkin mediated mitophagy [40] and basal degradation of MICOS [41]. In our previous studies, *SAMM50* KD recapitulated the phenotypes of *Chchd3* KD, resulting in reduced cardiac contractility and diminished sarcomeric actomyosin levels [3]. We therefore speculate that Chchd3 requires interaction with SAMM50 to regulate heart function and that loss of *Chchd3* likely disrupts this interaction, leading to perturbed mitophagy.

In addition, MICOS has been shown to localize in proximity of ER-mitochondria contact sites [42], which serve as hubs for the localized transfer of Ca^2+^ and ROS [43]. *Chchd3* KO in HeLa cells reduced the number of ER-mitochondria contacts and increased the distance between ER membrane and outer mitochondrial membrane [37]. While the elevated oxidative stress resulting from *Chchd3* KD may contribute to ER stress, it remains to be determined whether Chchd3 plays a direct role in maintaining ER-mitochondria contact site or mediating the transfer of Ca^2+^ and ROS.

In summary, our findings suggest that the three pathways, mitophagy, oxidative stress and ER stress, may be possible mediators and interactors in the MICOS genetic regulatory network.

## Material and Methods

### Fly stocks

Fly stocks from Bloomington Drosophila Stock Center, included w^1118^ control (#3605), attP2 control (#36303), Stinger (#84278), Chchd3/6 RNAiC1 (#51157), PINK1 RNAi (#55886), PINK1 OE (#51648), parkin RNAi (#31259), parkin OE (#51651), mito-QC (#91641), catalase RNAi (#34020), catalase OE (#24621), BiP OE (#5843), Ire1 RNAi (#35253), Xbp1 RNAiA (#25990), Xbp1 RNAiB (#36755), PERK RNAi (#35162), ATF4 RNAi (#25985) and ATF6 RNAi (#26211). The strong Chchd3/6 KD stock, Chchd3/6 RNAiA (#52251), was from Vienna Drosophila Resource Center. Xbp1 OE stock (#004794) was from FlyORF. The mouse spliced Xbp1 OE stock was a generous gift from Dr. Pedro Fernandez-Funez [31]. Hand^4.2^-Gal4 was a gift from Z. Han [23, 24].

Flies in genetic interaction studies were performed at 21°C or 25°C, as described in [3]. One-week-old female flies were used in all experiments. All animal experiments have been approved by the Institutional Animal Care and Use Committees (IACUC) at Sanford Burnham Prebys Medical Discovery Institute.

### Heart function analysis

For *in vivo* imaging and recording of heart function, the red fluorescence tdtK transgene was used [25]. Intact flies were anesthetized and adhered to a transparent coverslip and imaged by fluorescent Olympus BX63 microscope. Live recordings of heartbeat were captured by Hamamatsu C11440 ORCA-flash4.0LT digital camera and HCImage software. Analysis was automatically performed by an R-script [44] as in [26].

### ROS assay by DHE staining

We followed the procedure as described in [11]. Briefly, flies were anesthetized and dissected to expose the heart. Heart samples were quickly rinsed in PBS and treated in 1:1000 (v/v) DHE (Invitrogen D23107) solution diluted in ddH_2_O for 7-10 minutes at room temperature in the dark. Samples were washed in PBS for 3 times, 5 minutes each. After fixing with 7% formaldehyde for 5 minutes, heart samples were mounted in ProLong Gold Antifade Reagent with DAPI medium (Invitrogen P36935).

### Immunostaining

We followed procedures routinely used in the lab [3, 45]. Flies were anesthetized and dissected. Exposed heart samples were relaxed by 10mM EGTA, followed by fixation in 4% formaldehyde for 20 minutes. Heart samples were incubated in primary antibody overnight at 4°C. After washing out the primary antibody, samples were incubated in secondary antibody for 2 hours at room temperature. Samples were mounted in ProLong Gold Antifade Reagent with DAPI medium (Invitrogen P36935).

Primary antibodies included mouse monoclonal KDEL receptor antibody (1:100, Santa Cruz Biotechnology, sc57347), chicken IgY anti-GFP antibody (1:200, AVES, GFP1010), rabbit anti-mCherry antibody (1:200, Rockland, 600401p16) and anti-ATP5A (1:100, Abcam 14748) or anti-ATP5A1 (1:200, Invitrogen 43–9800). Secondary antibodies included Alexa Fluor 488 goat anti-chicken IgY (1:200, Jackson ImmunoResearch Labs, 103545155), DyLight 649 goat anti-mouse (1:200, ThermoFisher, 115-495-003), and Alexa Fluor 594 goat anti-rabbit 594 (1:200, Jackson ImmunoResearch Labs, 111585003).

### In-situ hybridization

Gene expression in adult hearts was assessed for BiP using HCR (Molecular Instruments) following the protocol developed by Bruce et al. [46] with slight modifications, as previously described [45]. Flies were dissected in oxygenated artificial hemolymph to expose the linear tube-like heart; excess fat was suctioned off with a micropipette, and hearts were fixed for 20 minutes in 4% paraformaldehyde. Hybridization and fluorescent labeling were performed according to manufacturers’ protocols.

All the images were observed by fluorescence microscope (BZ-X800, Keyence, Osaka, JAPAN) using 40X magnification.

### qRT-PCR

qRT-PCR was performed as in [45]. Briefly, for each biological replicate, 20-30 heart samples were collected from one-week-old female flies in either control or *Chchd3* KD group. The samples were homogenized for RNA extraction with miRNeasy Mini Kit (Qiagen 217004) and reverse transcription with QuantiTect Reverse Transcription Kit (Qiagen 205313). qPCR was performed with SYBR Green (Roche 04707516001) on a Roche LightCycler 96. The reference gene was rp49. The primer sequences used were 5’-AAACGCGGTTCTGCATGAG-3’ (rp 49 forward), 5’-GCCACCAGTCGGATCGATAT-3’ (rp49 reverse), 5’-GCTCAACCTGGATCTATTCC-3’ (BiP forward) and 5’-TGGATGGTGACGGTGTGCTGGTTAT-3’ (BiP reverse).

### Generation of human cardiomyocytes

Cardiomyocytes were generated as previously described [47] from Id1-overexpressing hiPSCs derived from dermal fibroblasts and donated by the laboratory of Dr Joseph Wu (Stanford University, CA, USA). HiPSCs were dissociated with 0.5 mM EDTA (Invirtogen, AM9260G) in PBS without CaCl_2_ and MgCl_2_ (Corning, MT21040CV) at for 37°C 7 min. hiPSCs were resuspended in mTeSR-Plus medium (StemCell Technologies, 100-0276) supplemented with 2 µM thiazovivin (StemCell Technologies, 72254) and plated in a Matrigel-coated 12-well plate (3×10^5^ cells per well) and cultured for 2 days until they reached ≥90% confluence to begin cardiomyocyte differentiation. Cells were treated with 6 µM CHIR99021 (Selleck Chemicals, S1263) in S12 medium for 48h followed by 2 µM Wnt-C59 (Selleck Chemicals, S7035) in S12 medium for 48h. Cells were cultured in fresh S12 media for a further 24h before being dissociated with TrypLE Express (Gibco, 12604013) for 2 min and blocked with RPMI 1640 medium (Gibco, 11875093) supplemented with 10% fetal bovine serum (FBS; Omega Scientific, FB-01) and being replated at 9×10^5^ cells per well) in S12 medium supplemented with 4 mg/l Recombinant Human Insulin (Gibco, 12585014) (S12+ medium), and 2 µM thiazovivin. The S12+ medium was refreshed after 36h and replaced after 48h of culture with Albumax media containing RPMI (Gibco) supplemented with 213 µg/µl L-ascorbic acid (Sigma-Aldrich), 500 mg/l BSA-FV (Gibco), 0.5 mM L-carnitine (Sigma-Aldrich) and 8 g/l AlbuMAX Lipid-Rich BSA (Gibco). After 5 days of culture in Albumax, cells were purified with lactate medium [RPMI without glucose, 213 µg/µl L-ascorbic acid, 500 mg/L BSA-FV and 8 mM sodium-DL-lactate (Sigma-Aldrich)], for 4 days before being switched back to Albumax medium for a further 6 days of culture.

### siRNA Knockdown, immunostaining and imaging of cardiomyocytes

hiPSC-derived cardiomycoytes at day 25 of differentiation were dissociated with TrypLE Select 10X (Gibco, 12605010) for 8 min, and TrypLE Select 10X was neutralized by adding RPMI supplemented with 10% FBS. Cells were resuspended in Albumax media supplemented with 2 µM thiazovivin and plated in 100µl at a density of 8000 cells per well in a Matrigel-coated 384-well plate containing 10µl of Opti-MEM (Gibco, 31985062) supplemented with 1% lipofectamine RNAiMAX (Invitrogen, 13778100) and 5µl siRNA (Dharmacon) at 0.5µM (25nM final concentration). Cells were cultured for 3 days to day 28 of differentiation. Prior to fixation, cells were treated with 10µM dihydroethidium (Invitrogen, D23107) for 30 minutes at 37°C. Cells were then fixed with 4% PFA for 10 minutes followed by permeabilized for 10 minutes with 0.1% Triton X in PBS and washed four times with PBS. Cells were then treated with blocking buffer for 1 hour. Cells were stained with TNNT2 (rabbit; Sigma-Aldrich, 017888), and KDEL (mouse; Santa Cruz, sc-57347) primary antibody in blocking buffer overnight at 4℃, then were washed 6 times with PBS at room temperature before being stained with secondary antibodies (donkey-anti rabbit 568 [A10042] and donkey-anti mouse 488 [A21202] and DAPI for 1h at room temperature, then washed four times with PBS. Cells were imaged using an ImageXpress Micro XLS microscope (Molecular devises) at 20X. Images were analyzed using INCarta software (Molecular devises) using a pretrained SINAP module to accurately segment DHE staining and a robust puncta algorithm was used for the detection of KDEL staining.

### Statistics

In all the functional analysis of fly heart, the bar graphs showed mean with SEM. Data were analyzed by one-way ANOVA with Bonferroni multiple comparisons. Significance was set as *<0.05, **<0.01, ***<0.001 and ns (not significant). In each panel, inside of the last data column at the bottom is indicated the interaction p-value from two-way ANOVA analysis. Statistical analysis of human cardiomyocyte data was performed using a one-Way ANOVA followed by a Dunn’s post-hoc test.

## Acknowledgements

We would like to thank Reed Walchle and Keyence for assistance in image acquisition and quantification.

## Figures

**Supplemental Figure 1:**
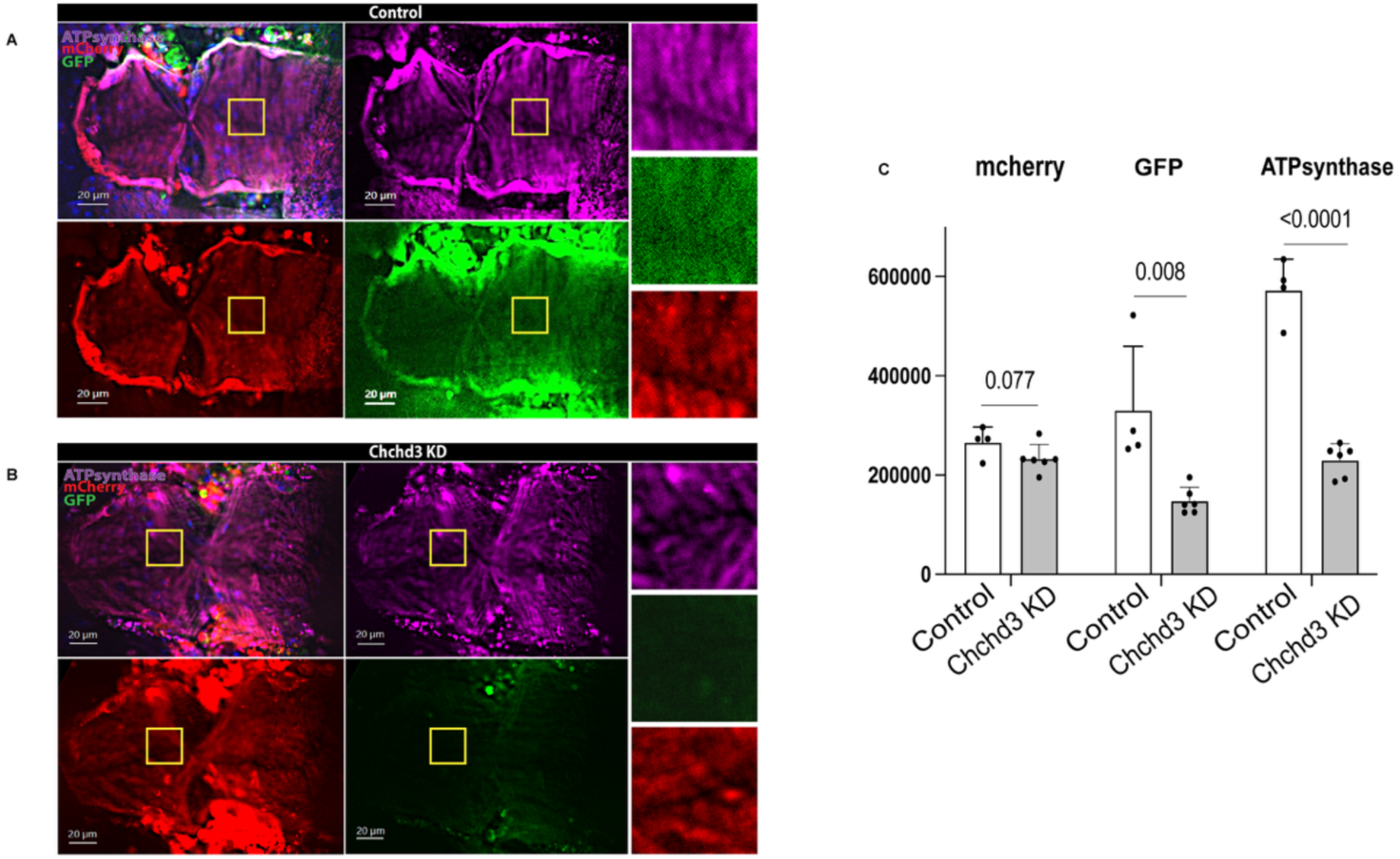
Increased degradation of dysfunctional mitochondria in *Chchd3* KD hearts. (**A,B**) Mitophagy was visualized by the mitochondrial-targeted mito-GFP-mCherry reporter and ATP synthase staining. Wildtype hearts exhibit robust GFP (green), mCherry (red) and ATP synthase (purple) immunofluorescence of intact mitochondria in the heart (**A**), whereas *Hand^4.2^*-Gal4-driven *Chchd3* KD hearts exhibit reduced in GFP and ATP synthase fluorescence, indicating degradation of mitochondria in lysosomes (**B**). Bar: 20 μm. Yellow squares indicate magnified areas on the right. **(C)** Quantification of mCherry, GFP and ATP synthase fluorescence intensity in wildtype and *Chchd3* KD hearts.

**Supplemental Figure 2.**
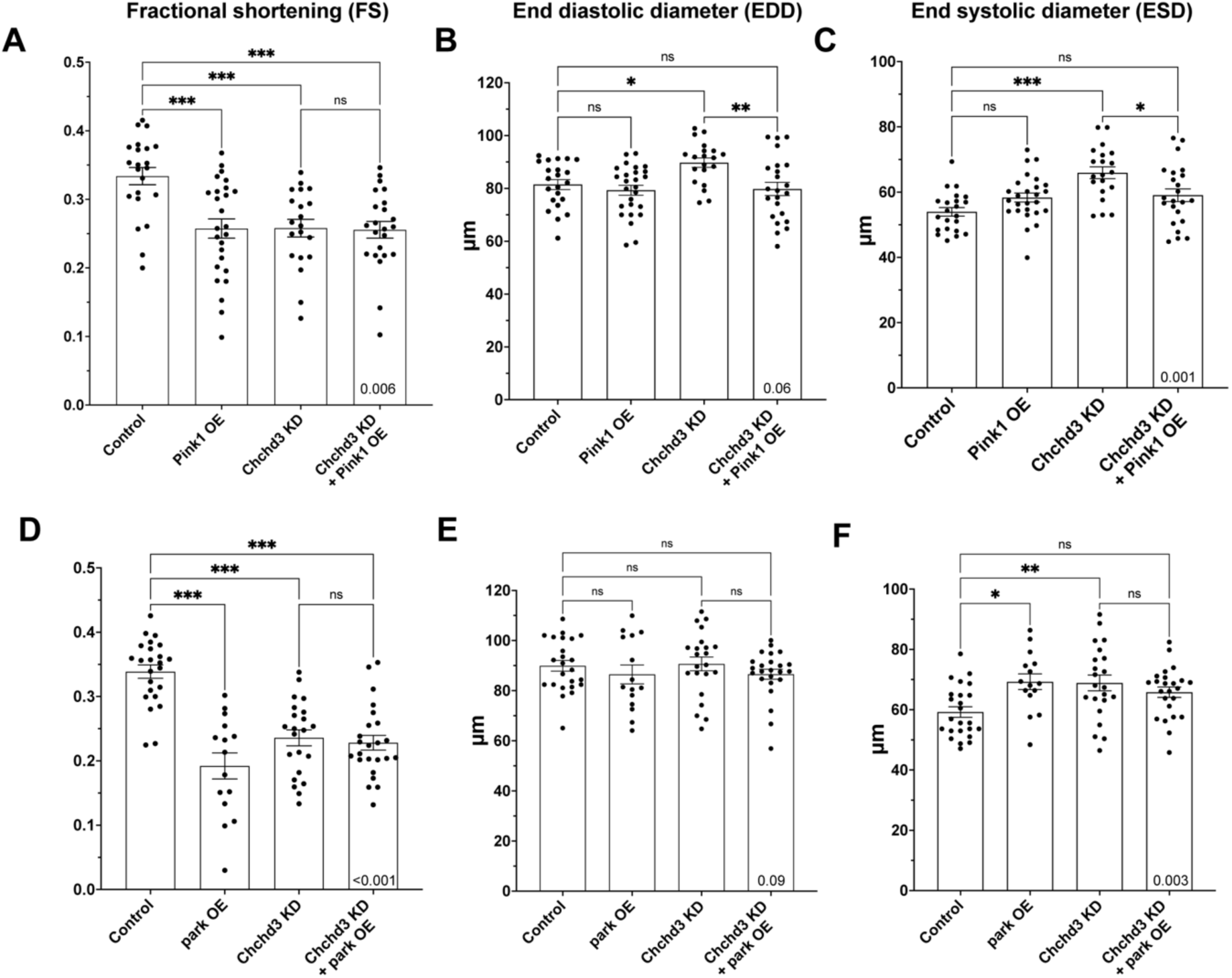
Promoting mitophagy by *Pink1* or *park* OE did not rescue the heart defects due to *Chchd3* KD. *Chchd3* KD at 25°C caused reduced heart contractility (**A,D**) and increased dilation (**B,C,E,F**). However, an attempt to promote mitophagy by *Pink1* or *park* OE did not result in rescuing contractility or diameters, except for a partial rescue by *Pink1* OE in EDD (**B**) and ESD (**C**) towards the control level. In fact, surprisingly, OE of *Pink1* or *park* by themselves had a detrimental effect on contractility (A, D).

**Supplemental Figure 3.**
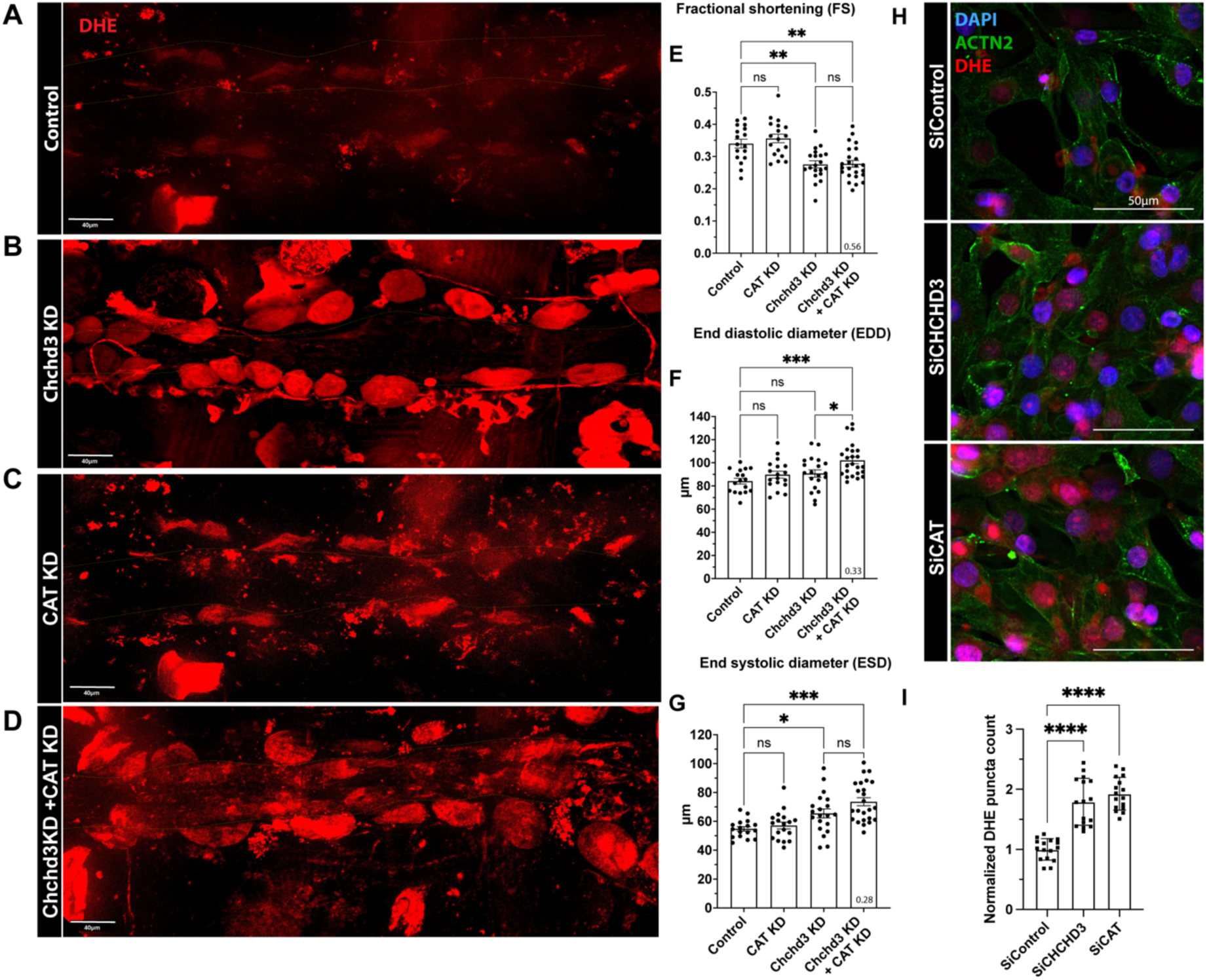
**(A-D)** ROS level examined by DHE staining of flies raised at 21°C. Compared to control (**A**), *Chchd3* KD (**B**), *CAT* KD (**C**) or in combination (**D**) increased ROS levels in cardiomyocytes and pericardial cells. (**E-G**) Increasing ROS levels by *CAT* KD did not further aggravate the compromised heart contractility due to *Chchd3* KD (**E**) but induced a moderate increase in diameters (**F-G**).

**Supplemental Figure 4.**
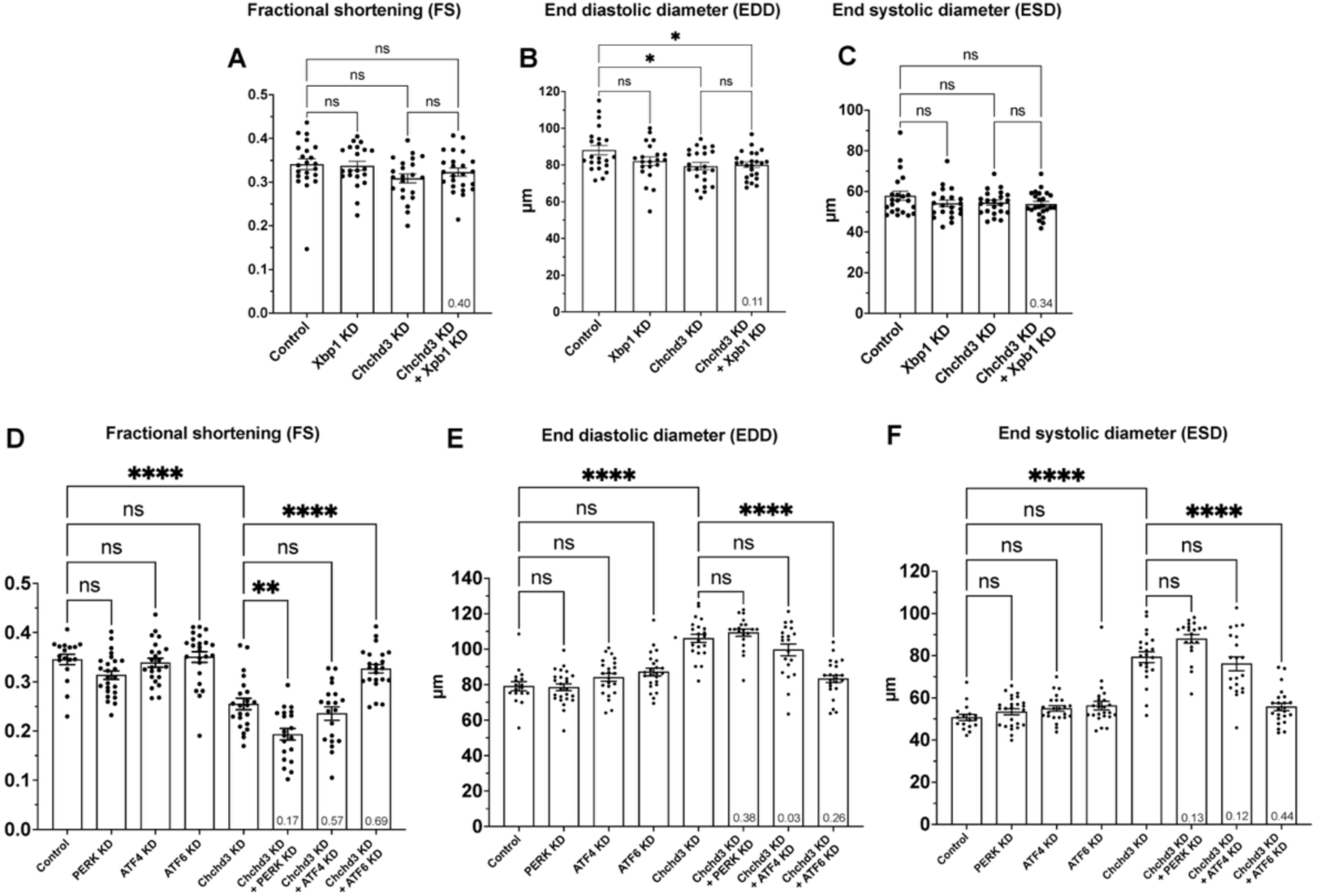
PERK-ATF4 and ATF6 branches of ER stress did not significantly interact with *Chchd3*. **(A-C)** *Xbp1* KD alone or in combination with *Chchd3* KD, at 21°C, did not exhibit any appreciable heart defects. **(D-F)** In contrast to ATF4 and ATF6 KD, combined KD at 21°C of *Chchd3* and *PERK* moderately aggravated cardiac defects, compared to *Chchd3* KD alone.

